# Errors in Predicting Muscle Fiber Lengths from Joint Kinematics Point to the Need to Include Tendon Tension in Computational Neuromuscular Models

**DOI:** 10.1101/2020.07.08.194381

**Authors:** Daniel A Hagen, Francisco J Valero-Cuevas

**Affiliations:** Department of Biomedical Engineering, University of Southern California, Los Angeles, CA, USA; Division of Biokinesiology and Physical Therapy, University of Southern California, Los Angeles, CA, USA

**Author notes:** Corresponding author, **University of Southern California**, *Department of Biomedical Engineering*, Brain-Body Dynamics Lab, Ronald Tutor Hall, RTH-421, 3710 S. McClintock Ave, Los Angeles, CA 90089-2905, USA, Website: ValeroLab.org. All authors were fully involved in the study and preparation of this preprint manuscript.

**Keywords:** kinematics, muscle fiber length, tendon dynamics, musculotendon, muscle mechanics, musculoskeletal modelling

## Abstract

Accurate predictions of tendon forces must consider musculotendon mechanics; specifically muscle fiber lengths and velocities. These are either predicted explicitly by simulating musculoskeletal dynamics or approximated from measured limb kinematics. The latter is complicated by the fact that tendon lengths and pennation angles vary with both limb kinematics *and* tendon tension. We now derive the error in kinematically-approximated muscle fiber lengths as a general equation of muscle geometry and tendon tension. This enables researchers to objectively evaluate this error’s significance—which can reach ~ 80% of the optimal muscle fiber length—with respect to the scientific or clinical question being asked. Although this equation provides a detailed functional relationship between muscle fiber lengths, joint kinematics and tendon tension, the parameters used to characterize musculotendon architecture are *subject*- and *muscle*-specific. This parametric uncertainty limits the accuracy of *any* generic musculoskeletal model that hopes to explain subject-specific phenomena. Nevertheless, the existence of such a functional relationship has profound implications to biological proprioception. These results strongly suggest that tendon tension information (from Golgi tendon organs) is likely integrated with muscle fiber length information (from muscle spindles) at the spinal cord to produce useful estimates of limb configuration to enable effective control of movement.

## 1. Introduction

Musculotendon (MT) complexes, as the name suggests, are composed of both muscle fibers *and* tendons [3]. These three-dimensional (often overlapping) bundles of fibers bulge, twist, and change lengths during contractions, but are conceptualized for simplicity as a flattened parallel bundle of fibers in series with an elastic element. This *parallelogram* simplification states that all fibers act in parallel *but* askew from the line of action by some pennation angle (*ρ*). This compartmentalizes the contributions of fiber length (*l_m_*) and tendon length (*l_T_*) to MT behavior (figure 1) [1–3]. Thus, MT length (*l_MT_*) is defined as the sum of *l_T_* (combining the tendons of origin and insertion; *l_T_* = *l*_*T*,1_ + *l*_*T*,2_) and the portion of *l_m_* projected onto the line-of-action of the MT (figure 1A, equation (1)) [3].

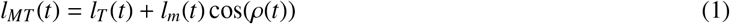

**Figure 1:**
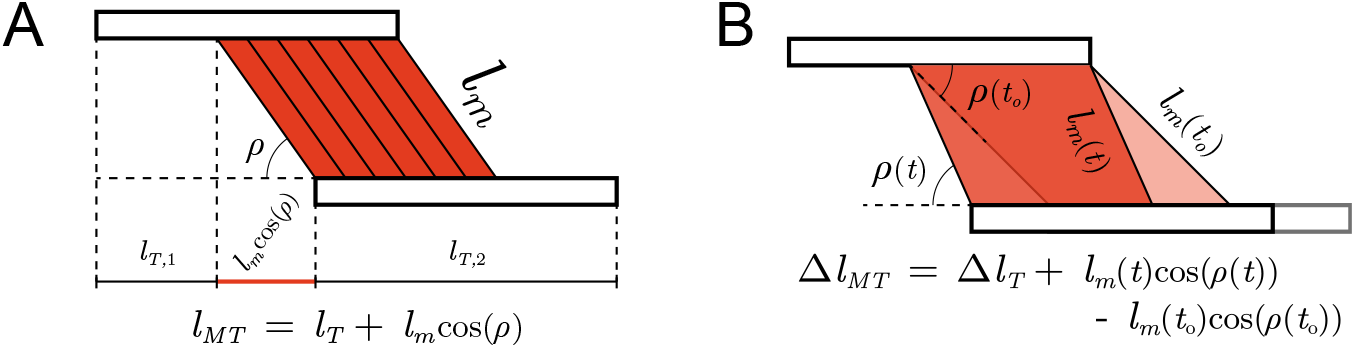
Approximation of musculotendon (MT) geometry as a flattened parallel bundle of pennated fibers in series with tendon (A) such that the change in length or *excursion* (Δ*l_MT_*, B) is defined as the sum of tendon length change (combining the tendons of origin and insertion, or Δ*l_T_* = Δ*l_T,1_* + Δ*l_T,2_*) and change in the portion of fiber length projected onto the line of action of the MT (*l_m_*(*t*) cos(*ρ*(*t*)) – *l_m_*(*t_o_*)cos(*ρ*(*t_o_*))) [1–3].

The excursion of the MT (Δ*l_MT_*, equation (2)) at time *t* is then defined as the difference between equation (1) evaluated at times *t* and *t_o_*, such that *l_m_* at time *t* is given by equation (3).

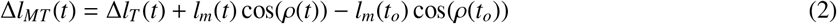

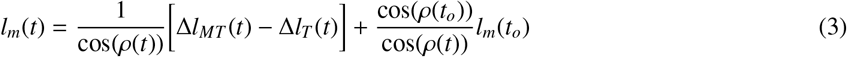

In practice, MT excursion can be calculated from the measured kinematics of a movement^1^. Even though *l_m_, l_T_*, and *ρ* are difficult to measure *in vivo* for a given subject, databases from imaging or cadaver studies can be used as approximations [4, 8–15]. This makes kinematics-based approximations for fiber lengths the most practical and most commonly used. These approximations further rely on the fundamental assumptions that tendons are inextensible (i.e., Δ*l_T_*(*t*) ≈ 0) or that *ρ* is either constant (*ρ*(*t*) ≈ *ρ_c_*) or negligible (*ρ*(*t*) ≈ 0).

When *only* assuming inextensible tendons (*IT*), equation (3) simplifies to:

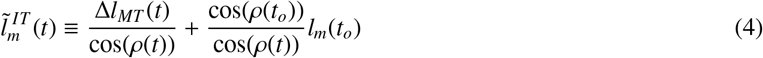

Alternatively, in the case where tendon stretch is included but *ρ* is assumed *constant* (*CP*), equation (3) simplifies to:

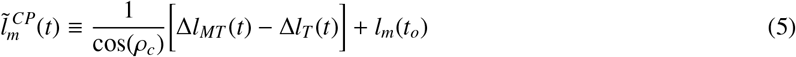

More often, however, both assumptions are made and the kinematics-based approximation to fiber length simplifies equation (3) to:

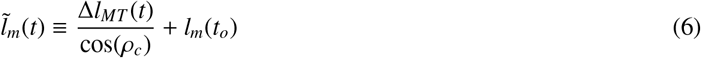

which, in effect, has led the field to often equate fiber length changes to MT excursions for small pennation angles [16]^2^.

Such approximations have greatly facilitated and enabled inferences regarding fiber lengths, velocities, or spindle (afferent) activity for multiple tasks [17–20]. Alternatively, the neuromechanical simulation software *OpenSim* offers a “stiff tendon” mode that makes similar assumptions [21]. In the aforementioned approaches, pennation angle is generally ignored or assumed constant. This practical approach to MT function has enabled computational studies to infer the neural control strategies of musculoskeletal systems [3, 16, 22–24], even though such models of muscle are known to be prone to parameter sensitivity [25, 26].

Here we explore how these approximations and sensitivities affect the conclusions that can be drawn from such muscle models—independently of the assumed contractile element. It is important to note that our analysis has implications to most lumped-parameter models (e.g., Hill-type models, [27]) and population-of-muscle-fibers models (e.g., the Fuglevand model, [28]) because such models tend to ignore tendon mechanics and consider pennation angles to be negligible. We first explicitly derive equations for the errors produced when making a variety of assumptions to enable the reader to make informed decisions about their impact on the particular scientific question or clinical condition studied. We then go on to explore the consequences to inter-muscle and inter-subject variability, and its impact on studies of neuromuscular control.

## 2. Methods

### 2.1. Derivation of Tendon Deformation as a Function of Tendon Forces

The elastic properties of collagen are such that tendon force (*f_T_*) depends on tendon length (*l_T_*), with a characteristic nonlinear “toe” region at low forces followed by a linear region at higher forces [3, 29–31]. To generalize this relationship across muscles, [31] modelled the *normalized* tendon force-length curve (equation (7a)) whereby *f_T_* was normalized by the maximum isometric force *of the muscle* 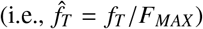 and *l_T_* was normalized by the *optimal* tendon length 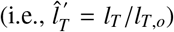. Additionally, the parameters *c^T^*, *k^T^*, and 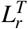 fit the asymptotic slope, curvature, and lateral-shift, respectively^3^. Note that [31] decided to normalize *l_T_* by the *optimal* tendon length (*l_T,o_* — the tendon length when the muscle produces its maximal isometric force) instead of the slack length (*l_T,s_* – tendon length when tendon force is negligible) as it produced more “congruent curves.” However, the literature reports the ratio between tendon slack length and optimal muscle length [3, 32], so we preferred to normalize *l_T_* by the slack length. The same relationship can be rewritten as equation (7b) for when *l_T_* is normalized by *l_T,s_* instead 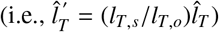.

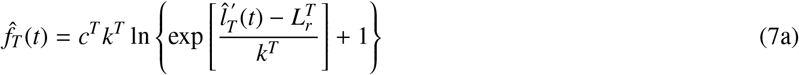

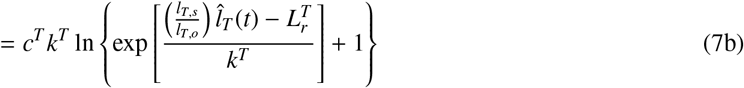

This relationship, albeit *muscle*- and *subject*-specific, can be inverted to provide tendon length as a function of tendon force (equation (8))^4^.

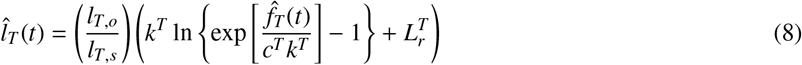

Therefore, the normalized change in tendon length can be rewritten as a function of both the current and initial forces on the tendon (equation (9)).

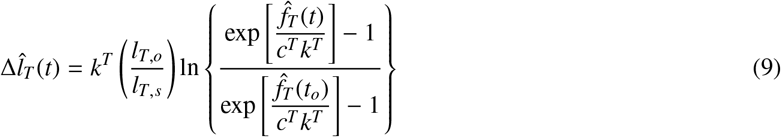

### 2.2. Error in Fiber Length Approximations

To allow for better comparison across muscles, we define the relative errors in kinematically-approximated fiber lengths (denoted by *η*) as the differences between equation (3) and equations (4)–(6), normalized by optimal fiber length (*l_m,o_*). When assuming only *IT*, the error is defined by equation (10). Intuitively, this error will be equal to the tendon length change that was ignored (equation (9)), projected back onto the line of action of the fibers

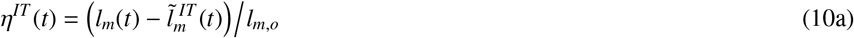

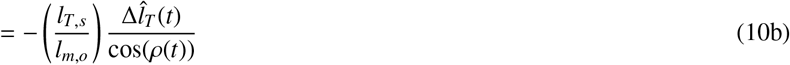

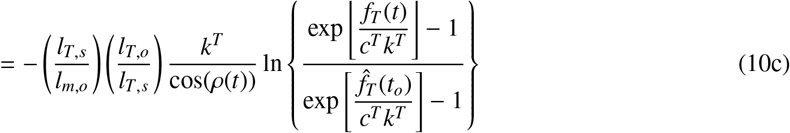

Alternatively, when including tendon length changes but assuming *CP* the error is defined by equation (11).

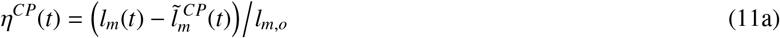

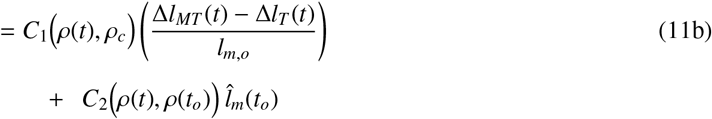

Where *C*_1_ reflects the proportion of Δ*l_MT_* − Δ*l_T_* that was not projected back onto the line of action of the fibers but instead some other axis given by *ρ_c_* (equation (12)) and *C*_2_ represents the proportion of *l_m_*(*t_o_*) incorrectly projected back onto the current line of action of the fibers (equation (13)).

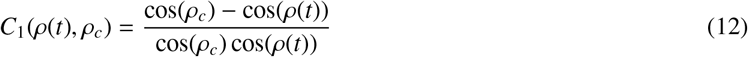

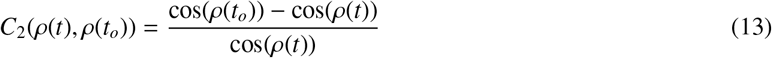

These now allow the relative error in kinematically-approximated fiber lengths, *η*(*t*), to be defined as equation (14).

Note that this error accounts for the proportions of Δ*l_MT_* and *l_m_*(*t_o_*) that were not mapped onto the line of action of the fibers (*CP* assumption) as well as the ignored tendon length change (*IT* assumption).

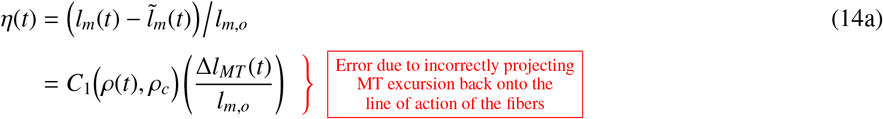

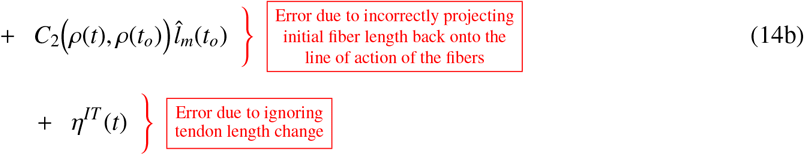

Finally, correcting for this error provides a more accurate approximation of normalized fiber length that takes both limb kinematics and tendon tension in account (equation (15)).

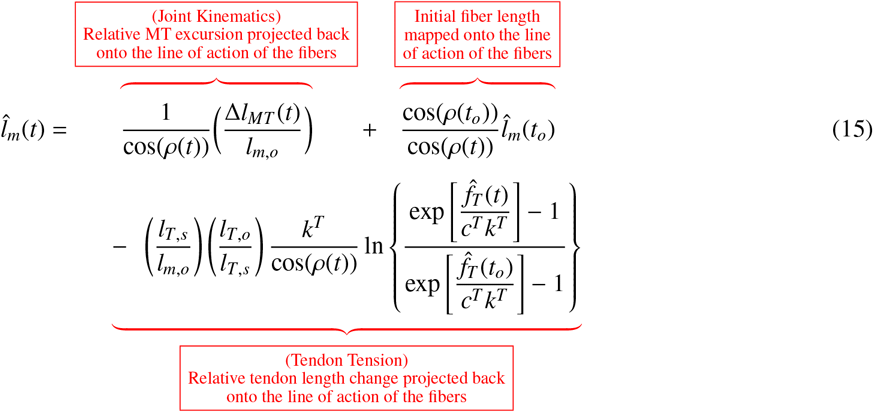

## 3. Results

### 3.1. Error from Ignoring Fiber Pennation

As the first two terms of (14b) illustrate, assuming that pennation angle is either constant or negligible will result in errors when approximating fiber lengths from limb kinematics [33]. The coefficient of the first term (*C*_1_) reflects the percentage of Δ*l_MT_* that was not projected back onto the line of action of the fibers, but instead some other axis given by *ρ_c_* (equation (12), figure 2). While this error will be negligible when *ρ*(*t*) ≈ *ρ_c_*, its is more sensitive to small changes in pennation as *ρ_c_* increases (figure 3A)^5^. Therefore, when assuming CP for muscles with larger pennation angles, the same deviation (*ρ* = *ρ_c_* ± *δρ*_1_) will result in a larger percentage of Δ*l_MT_* that was incorrectly projected back onto the line of action of the fibers.

**Figure 2:**
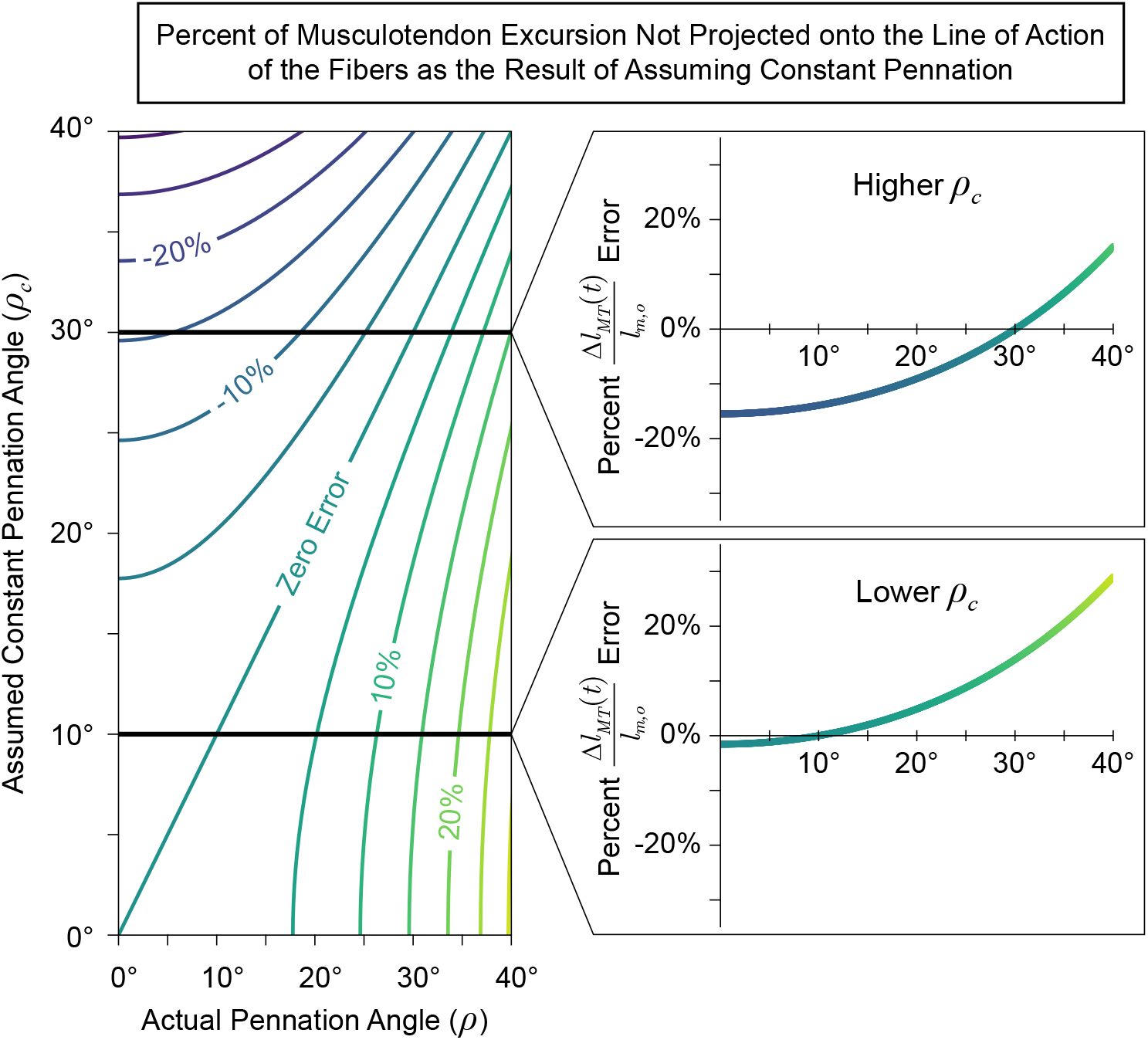
Contour map (*left*) for the percentage of musculotendon excursion that would be incorrectly mapped onto the line of action of the fibers (C_1_) as the result of assuming some constant pennation angle as a function of the true pennation angle. For any assumed value of constant pennation (*ρ_c_*), the resulting plot of *C*_1_ is given by the corresponding horizontal cross-section of the contour plot. Examples of plots for lower and higher values of *ρ_c_* are shown on the *right*. Note that the error is negligible when the true pennation angle is equal to the assumed value (diagonal line on the *left*, and zero-crossings on the *right*). Additionally, the error is less than ±5% when the assumed *and* actual pennation angles are less that ~ 18°.

**Figure 3:**
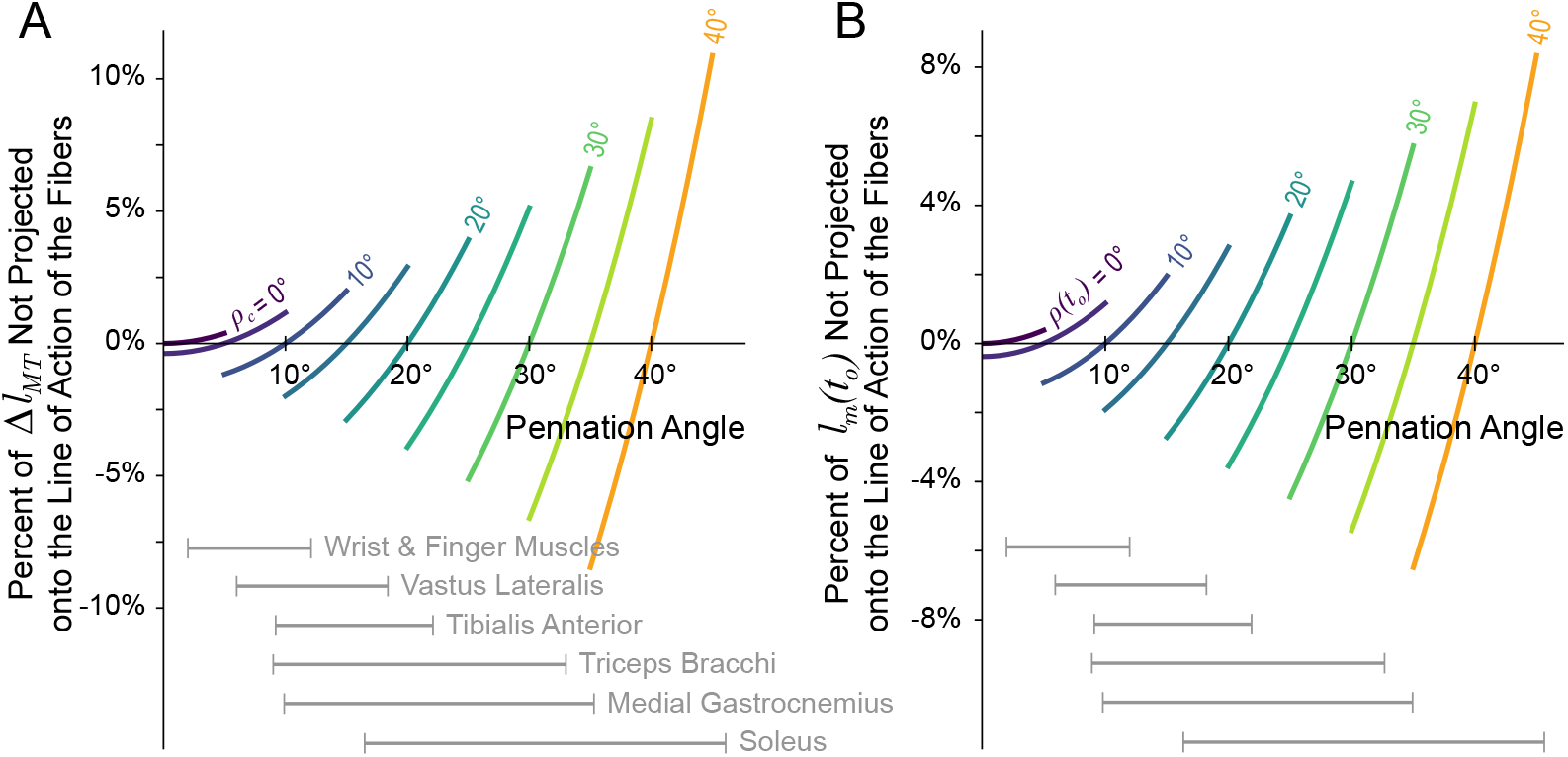
Sensitivity of the relative error coefficients *C*_1_ (the proportion of MT excursion not projected back onto the line of action of the fibers, A) and *C*_2_ (the proportion of the initial fiber length not projected back onto the line of action of the fibers, B) from equation (14). A small deviation (±5°) was applied to *ρ_c_* or *ρ*(*t_o_*), respectively, and the resulting change in the coefficients were plotted. For (A), while the error is minimized when *ρ*(*t*) = *ρ_c_*, as *ρ_c_* increases the same deviation from the true pennation angle (*ρ*(*t*) = *ρ_c_* ± 5°) will produce larger changes in *C*_1_ and, therefore, a larger percentage of Δ*l_MT_* would be incorrectly projected back onto the line of action of the fibers. Similarly for (B), the error will be minimized when the pennation angle does not change from the initial value (i.e., *ρ*(*t*) = *ρ*(*t_o_*)), but as *ρ*(*t_o_*) increases, the same deviation from the initial value (*ρ*(*t*) = *ρ*(*t_o_*) ± 5°) will result in larger changes to *C*_2_ and, therefore, a larger proportions of the initial fiber length would be incorrectly mapped back onto the fibers at time *t*. Ranges of pennation angles reported in the literature have been provide for a few muscle groups for reference [10, 14, 34–39].

The coefficient of the second term (*C*_2_) reflects the percentage of *l_m_*(*t_o_*) incorrectly projected back onto the line of action of the fibers as the result of assuming constant pennation (figure 4). Similar to *C*_1_, this error will be negligible when the pennation angle does not deviate from the initial value (*ρ*(*t*) ≈ *ρ*(*t_o_*)) and becomes more sensitive to changes in pennation as *ρ*(*t_o_*) increases (figure 3B)^6^. Therefore, when assuming CP for muscles with larger pennation angles that *also undergo larger pennation angle changes*, the same deviation (*ρ*(*t*) = *ρ*(*t_o_*) ± *δρ*_2_) will result in a larger percentage of *l_m_*(*t_o_*) that was not projected back onto line of action of the fibers.

**Figure 4:**
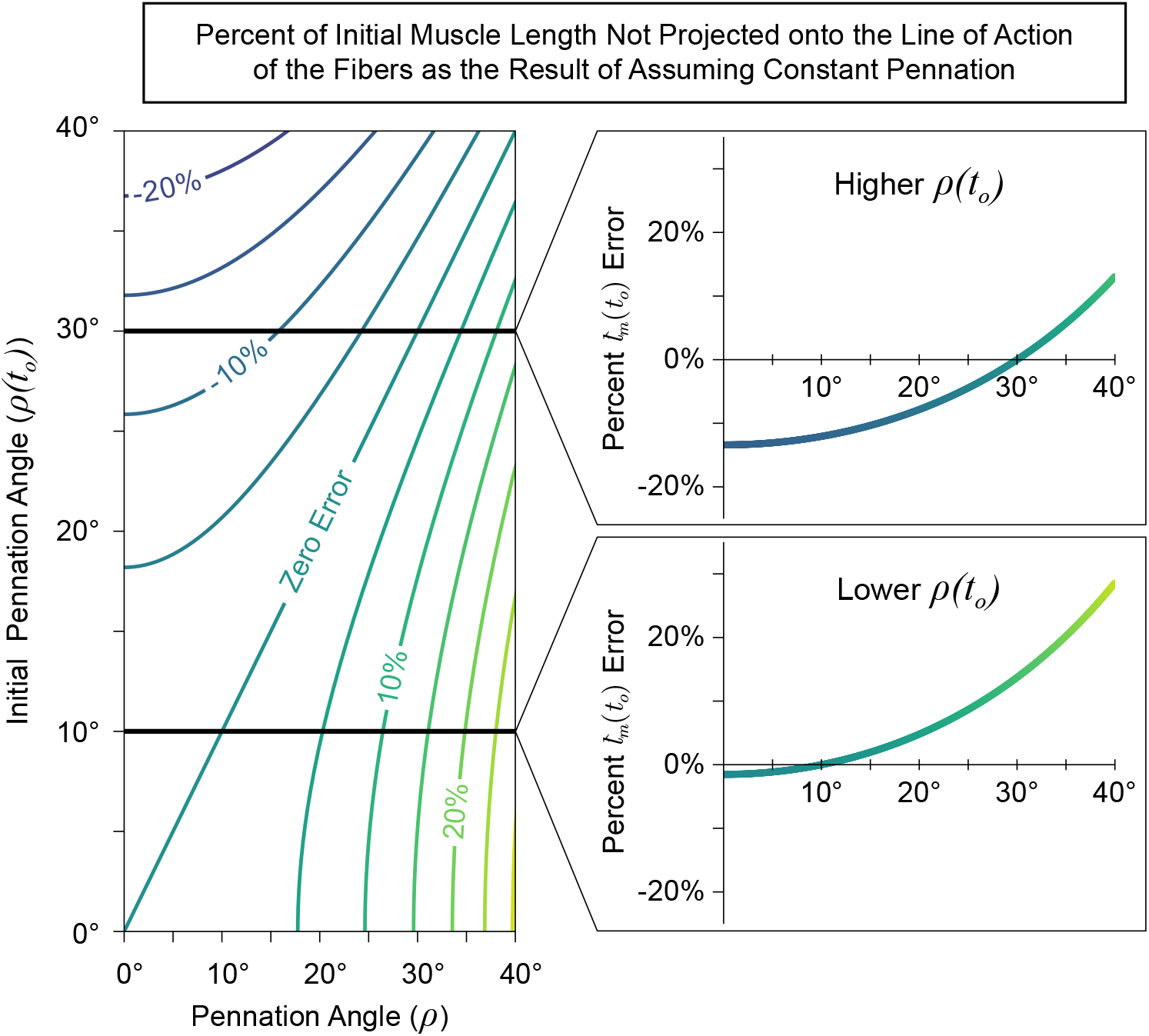
Contour map (*left*) for the percentage of initial fiber length that would be incorrectly mapped back onto the true line of action of the fibers (*C*_2_) as the result of assuming some constant pennation angle as a function of the initial and current pennation angles. Regardless of the assumed constant pennation value, this error will depend on the amount by which the pennation angle changes from its initial value. Therefore, for some initial pennation angle, the resulting plot of the coefficient *C*_2_ is given by the corresponding horizontal cross-section of the contour plot. Examples of plots for lower and higher values of *ρ*(*t_o_*) are shown on the *right*. Note that the error is negligible when the current pennation angle is equal to the initial pennation angle (diagonal line on the *left*, and zero-crossings on the *right*). Additionally, the error is less than ±5% when the initial *and* actual pennation angles are less that ~ 18°. Therefore, for muscles with small pennation angles (< 18°) that *do not drastically change* over the course of a movement, this error will be relatively small (< ±5%).

Reported average pennation angles vary across muscles in humans; from ~ 1.3° to ~ 46.1° in the lower limb [10, 34–36] and from ~ 2.0° to ~ 33.0° in the upper limb [14, 37–39] with large variability reported across subjects. Additionally, pennation angles have been reported to change by 120 to 175% from rest when the muscle is fully activated [38, 40–44]. Therefore, for some muscles and movements, the percentage of either Δ*l_MT_* or *l_m_*(*t_o_*) that can be excluded or incorrectly mapped back onto the line of action of the fibers can be quite large (~ ±20%). As an example, [44] reported that the *medial gastrocnemius* experienced a Δ*l_MT_* ≈ −2 cm during a maximum squat jump, where the pennation angle increased from 20° to 35°. If the pennation angle had been assumed to be constant and equal to the initial pennation angle (i.e., *ρ_c_* = *ρ*(*t_o_*) = 20°), then 15.7% of Δ*l_MT_* and 14.7% of *l_m_*(*t_o_*) would have been incorrectly mapped back onto the fiber, resulting in an normalized error of around −6.5% (−3.13mm) and 17.2% (8.24mm), respectively^7^.

### 3.2. Error from Assuming Inextensible Tendon

The third term of (14b) represents the error associated with assuming inextensible tendons [45]. Figure 5 shows equation (14b) evaluated at two initial tendon tensions. Note that when 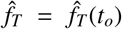 (i.e., the normalized tendon force intercept), the tendon will have undergone a net zero length change and the error will be zero. However, notice that the slope at the intercept (red lines) demonstrates that starting at lower forces (i.e., an intercept towards the left) creates greater sensitivity to deviations from the initial tension, whereas at higher forces the sensitivity is lower and approaches the asymptotic slope (black arrows). Therefore, tasks that simulate non-isotonic tendon forces at low levels (like most activities of daily living) are at the greatest risk of errors of this type. However, the parameters used to characterize the shape of the curves in figure 5 (i.e., *c^T^*, *k^T^*, *l_T,o_/l_T,s_*, and *l_T,s_/l_m,o_*) vary across muscles and subjects and can change with exercise, injury, or pathology [9, 46–52]. Additionally, *p* changes across muscles and subjects, as well as under different force levels for a given muscle (as previously stated) and will affect the error proportionally.

**Figure 5:**
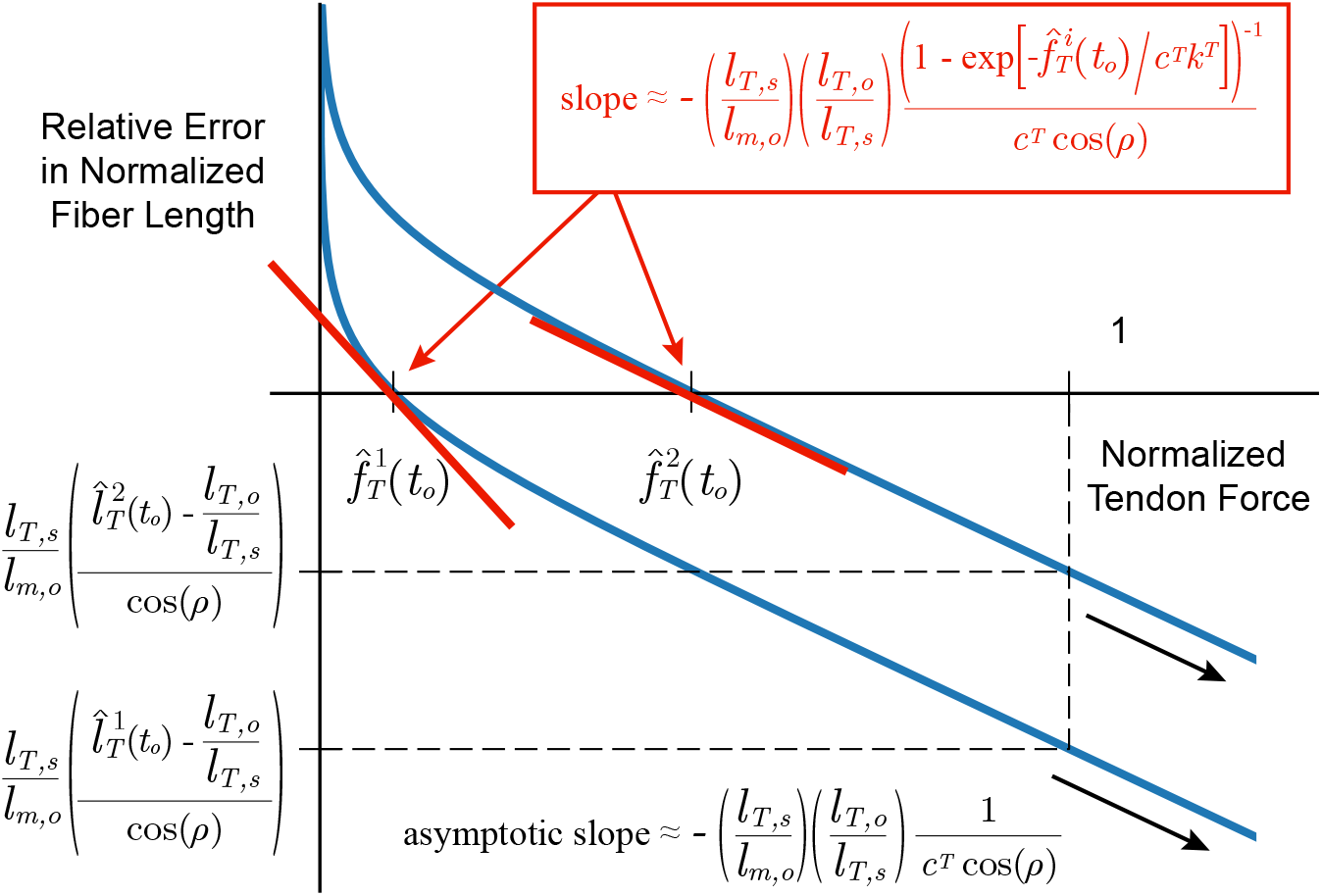
Relative error in fiber length as a result of assuming inextensible tendons (equation (10c)) that accounts for the previously ignored tendon length change, scaled by the tendon slack length to optimal fiber length ratio (*l_T,s_/l_m,o_*), and projected back onto the line of action of the fibers. Note that the error will be zero when the tension of the tendon is equal to the initial tension 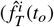—i.e., no net deformation of the tendon has occurred). Two different initial tension values have been chosen to demonstrate that starting at lower forces (i.e., an intercept towards the left) creates greater sensitivity to deviations from the initial tension, whereas at higher forces the sensitivity is lower and approaches the asymptotic slope (black arrows) where it will be proportional to *lτ_T,s_/l_m,o_* and inversely proportional to *c^T^* = *E* · *CSA_T_/F_MAX_* (i.e., the tendon’s normalized asymptotic stiffness).

Therefore, we explored how changes in these parameters affect the magnitude of the error in fiber lengths by simulating isometric force tasks from rest. By doing so, MT excursion will be zero such that kinematically-approximated fiber lengths will be constant and the subsequent error will be identical to the normalized length change of the tendon, scaled by *l_Ts_/l_m,o_* and divided by the cosine of the pennation angle. Figure figure 6A shows 10,000 random isometric force tasks (between 0-100% of maximum voluntary contraction, MVC) with the 5 parameters of interest uniformly sampled within their reported physiological ranges [53]^8^. Based on the reported range for *l_T,o_/l_T,s_* (and the definitions of *l_T,s_* and *l_T,o_*) the maximal normalized tendon deformation we can expect at MVC will be

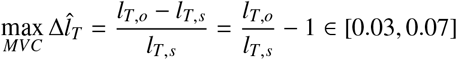

such that the maximal error in fiber length for this task will be the maximal tendon deformation scaled by the ratio *l_T,s_/l_m,o_* and divided by the cosine of the pennation angle for a given muscle. For physiological ranges of these values, the maximal error in fibers can reach magnitudes of ~80% *l_m,o_*. As expected, for higher tendon forces (≥ 75% of MVC) where 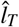 converges to *l_T,o_/l_T,s_*, those trials with larger errors (≥ 30% *l_m,o_*) shows no clear dependence on *c^T^* or *k^T^*, but instead are determined by the product of *l_T,o_/l_T,s_* and *l_T,s_/l_m,o_* (figure 6B). Surprisingly, for lower tendon forces (≤ 50% of MVC), errors in fiber lengths ≥ 30% (figure 6C) typically occur for musculotendons with *lower* normalized asymptotic tendon stiffness (*lower c^T^*), *lower* “toe” region curvature in the tendon’s force-length curve (*higher k^T^*), and higher ratios of tendon slack length to optimal muscle length (*l_T,s_/l_m,o_* > 6) and optimal tendon length to tendon slack length (*l_T,o_/l_T,s_* > 1.04). These parameters corresponds to a tendon that (i) is substantially longer than the muscle (i.e., larger *l_T,s_/l_m,o_*), (ii) can undergo relatively larger overall deformations (i.e., larger *l_T,o_/l_T,s_*), and (iii) has a *flatter* “toe” region in the tendon force-length curve. This “perfect storm” elicits disproportionately greater tendon deformations per unit force at lower forces. Conversely, if any of these conditions are not met, the errors in fiber lengths can be low (≤ 5% *l_m,o_*, figure 6A). Lastly, it is of course trivial to show from equations (2) & (3) that increasing the pennation angle will increase the proportion of Δ*l_T_* projected back onto the line of action of the fibers (i.e., a 1/ cos(*ρ*(*t*)) function), and thus increase the magnitude of this error. However, we see that even muscles with low pennation angles are not immune to high errors (see second column from right in figure 6B,C). To further explore the consequences of varying these parameters during an isometric task, the reader is pointed to the online interactive version of figure 6, available at https://daniel8hagen.com/images/tendon_length_change_parallel_coords.

**Figure 6:**
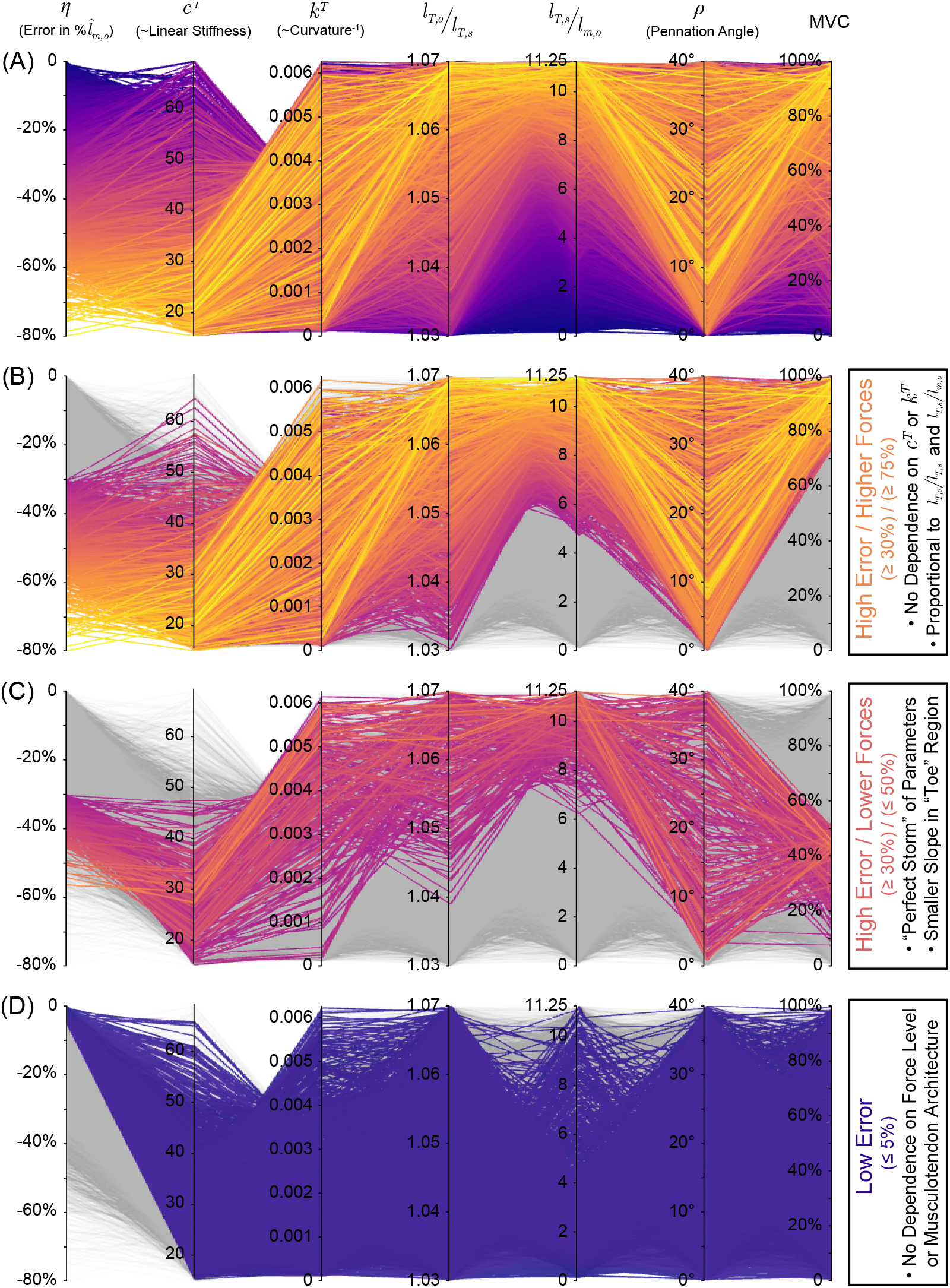
Parallel coordinates plot for the relative error in muscle fiber length (η) associated with assuming inextensible tendons for 10,000 random samples in the 5 parameters of interest (*c^T^, k^T^, l_T,o_/l_T,s_, l_T,s_/l_m,o_*, and *ρ*) within their reported physiological ranges during random isometric force tasks (*top*). As there is no MT excursion (Δ*l_MT_*) during an isometric contraction, assuming inextensible tendon is equivalent to assuming that muscle fiber length is constant at a given percentage of maximum voluntary contraction (MVC) and the error, therefore, will be identical to the normalized length change of the tendon, scaled by *l_T,s_/l_m,o_* and *l_T,o_/l_T,s_*, and divided by the cosine of the pennation angle. For higher forces (≥ 75% MVC), we find that high errors (≥ 30% *l_m,o_*) can occur for all values of *c^T^* and *k^T^* as, by definition, the tendon length converges to its optimal length (i.e., the error is only proportional to *l_T_,_o_/l_T,s_* and *l_T,s_/l_m,o_, secondfrom top*). Alternatively, we find that the error can be equally large for *lower* forces (≤ 50% MVC) when the tendon has *low* stiffness, *low* curvature (*high* radius of curvature), and *larger* ratios of *l_T,s_* to *l_m,o_* and *l_T,o_/l_T,s_* (i.e., the “perfect storm”, *second from bottom*). Conversely, if any of these conditions are not met, the errors in muscle fascicle lengths can be low (bottom). Lastly, pennation angles do not appear to preclude any muscles from this sort of error, but it is trivial to show that increasing ρ will increase the proportion of Δ*l_T_* projected back onto the line of action of the muscle fascicles. Vist https://daniel8hagen.com/images/tendon_length_change_parallel_coords to access this interactive parallel coordinate plot online.

## 4. Discussion

Biomechanics and neuromuscular control depend uniquely on the assumed properties of muscle. Chief among them is the dependence of muscle force for a given amount of neural drive on the lengths and velocities of its fibers. Our detailed analysis of how fiber lengths (and, by extension, velocities) are approximated from experimental data or simulations points to assumptions regarding musculotendon architecture (i.e., pennation angle and tendon elasticity) as the source of three dominant types of uncertainty and error (equation (14b)). More importantly, the magnitude of these errors is highly sensitive to the parameters characterizing musculotendon architecture. The significance of these potentially large errors is ultimately left to the modeller and their audience. To guide these decisions in practice, however, we provide a detailed presentation of the interactions between musculotendon architecture, musculotendon force magnitude, as well as its dependence on the ratios of tendon slack length to optimal fiber length and of optimal tendon length to tendon slack length. These high-dimensional interactions show that some combinations of parameters will produce (i) relatively low errors vs. (ii) a “perfect storm” where errors will be unacceptably large by any measure, even for relatively low force magnitudes. Our work highlights that extreme care must be taken when making such approximations in the context of the scientific question being asked, the muscles and tasks being studied, and the available experimental data.

We began by showing that, when approximating fiber lengths from the kinematics, separately assuming either constant pennation or inextensible tendons will incur distinct errors with respect to the ground truth. The magnitude of these errors are themselves highly sensitive to the parameters that characterize the musculotendon (i.e., fiber pennation angle and tendon elasticity).

It is intuitively obvious that assuming zero or constant pennation will incorrectly map the changes in musculotendon length and the initial fiber length onto the current line of action of the fibers. Our community has often assumed that this error is negligible for “small” pennation angles, which may be true (figure 3A). However, we show that starting at reasonably modest pennation angles of 20°, the sensitivity of the errors escalates exponentially to the point where a ±5° deviation from the assumed constant pennation angle can lead to > 5% of the musculotendon excursion to be unaccounted for in the fiber length approximation—with potentially important consequences to force-length and force-velocity calculations. Similarly, it is known that some muscles with modest pennation angles nevertheless undergo large changes in pennation during everyday movements [c. 120–175%, 38, 40–44]. In these cases, the amount of initial fiber length that is incorrectly mapped back onto the line of action of the fibers can be >> 4% as calculated for a ±5° change in pennation (figure 3B) and can reach magnitudes on the order of ~ 20% (see [44] example in Section 3.1). Both of these errors increase greatly for more pennate (and highly studied) muscles such as *tibialis anterior*, *triceps bracchi*, *medial gastrocnemius*, or *soleus*. This highlights the need for perhaps unrealistically or impractically accurate measurements of muscle pennation angle (and their change during a task) to accurately simulate force production in these muscles when they are operating away from the plateau of the force-length curve, and most everywhere in the force-velocity curve.

Additionally, if one wanted to assume inextensible tendons, one must note that the errors in estimated fiber lengths are exacerbated at lower forces (≤ 50% of MVC) if the combination of parameters that characterize the tendon’s normalized force-length relationship (i.e., its asymptotic stiffness, *c^T^*, and its curvature constant, *k^T^*) and the ratios of the tendon’s slack length to the optimal fiber length (*l_T,s_/l_m,o_*) and optimal tendon length to tendon slack length (*l_T,o_/l_T,s_*) meet the “perfect storm” criteria(e.g., *soleus* muscle) [3, 51, 55]. That is, errors can be quite large for more compliant tendons (i.e., low values of *c^T^* and high values of *k^T^*) with moderately high values of *l_T,s_/l_m,o_* and *l_T,o_/l_T,s_* (figure 6C). Conversely, if *any* of these “perfect storm” conditions are not met, the error can potentially be small, as can be seen by the wide distribution of parameters that produce less than 5% error in fiber lengths in figure 6D. Therefore, it is important to understand how a *particular* choice of parameters can affect the robustness of a kinematics-based fiber length approximation by understanding how they will effect the shape of the tendon’s force-length relationship and how deformations of tendon will subsequently produce scaled changes in fiber length.

The limitations of this study do not necessarily affect the validity of our results. For example, we did not consider more complex musculotendon architectures that do not reflect the simple parallelogram, and we ignored hysteresis, tendon creep, short-range stiffness, and force-relaxation. In fact, the added nonlinearities and state-dependence of these omissions suggest that we may, in fact, be underestimating these uncertainties and errors. By correcting for the errors explicitly derived here, we arrive at a more accurate approximation of fiber length that relies on both limb kinematics *and* tendon dynamics (equation (15)). This equation creates a functional relationship, through the limb kinematics, between fiber lengths and tendon tensions for concentric, eccentric and isometric contractions. However, in practice, its sensitivity to parameter values makes it all-but-impossible to use to recover specific musculotendon behavior without accurate *subject*- and *muscle-specific* parameters.

These results raise further important issues relevant to the role of muscle spindles to provide proprioception for neuromuscular control. In Nature, proprioception provides animals a robust awareness of the state of their body and of their relation to the environment. The muscle spindles embedded in the intrafusal fibers of muscle are known to provide information about the velocity and length of a muscle. Muscle spindles, by sensing along the line of action of the muscle fibers, are nevertheless subject to similar uncertainties and errors to those described above when estimating the length of the musculotendon. However, it stands to reason that the nervous system can learn to interpret and integrate spindle signals to estimate the posture of the limb because their afferent signals are implicitly functions of the *subject*- and *muscle-specific* musculotendon parameters. Nevertheless, equation (15) points to the physical necessity of knowing tendon tension in addition to fiber length to accurately estimate length (and velocity) of the musculotendon—and therefore limb posture. We speculate that this obligatory functional relationship between fiber lengths and tendon tensions may point to an additional evolutionary pressure for Golgi tendon organs (mechanoreceptors for tendon tension) whose projection to the same spinal, sub-cortical and cortical areas would critically enable more accurate estimates of limb posture.

## Data Accessibility

All data used in this paper were previously published.

## Author Contributions

D.A.H and F.J.V.-C. designed the research; D.A.H. performed the research; D.A.H. and F.J.V.-C. analysed the data. All authors discussed the results and wrote the paper.

## Competing Interests

We have no competing interests.

## Funding

Research reported in this publication was supported by the National Institute of Arthritis and Musculoskeletal and Skin Diseases of the National Institutes of Health under award numbers R01 AR-050520 and R01 AR-052345 and by the National Institute of Neurological Disorders and Stroke of the National Institutes of Health under the award number R21-NS113613 to F.J.V.-C. This work was also supported by Department of Defense CDMRP Grant MR150091 and Award W911NF1820264 from the DARPA-L2M programme to F.J.V.-C.

## Acknowledgements

We are grateful to Felix Zajac for inspiring this research and to Gerald Loeb for his seminal work on musculoskeletal systems and for his equation describing normalized tendon tension-deformation (derived in [31]) which allowed us to find an explicit form of this error. Additionally, we thank Akira Nagamori and Christopher Laine for their helpful feedback during early project development.

## Disclaimer

The content of this endeavour is solely the responsibility of the authors and does not represent the official views of the National Institutes of Health or the Department of Defense.

## A. Appendix

## A.1. New and Improved Equations for Musculotendon Length and Velocity

When simulating muscle fiber and tendon behavior it is paramount to use accurate equations for musculotendon (MT) length (*l_MT_*) and velocity (*v_MT_*). Historically, *l_MT_* has been calculated either by tracing the MT routing from origin to insertion across postures [4], or by approximating the MT excursion (*s*) induced by joint rotations at every joint crossed [5–7]. The latter approach assumes constant moment arms such that the MT excursion produced by a joint’s rotation from neutral (*θ* − *θ_o_*) would be equal to the arc length of a sector of a circle with radius equal to the moment arm (r, equation (18)). Therefore, *l_MT_* is approximated as the sum of some neutral MT length (*l_MTo_*) and the MT excursions of all joints crossed (equation (19)). Note that the negative sign in the MT excursion equation ensures that for a positive joint rotation from neutral, a MT with a positive moment arm will shorten.

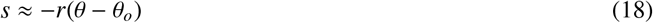

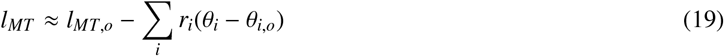

Musculotendon velocity (*v_MT_*) is defined as the change in excursion due to joint rotation(s) over time (20a). For constant moment arms, this is approximated as the linear combination of joint velocities 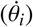 scaled by their moment arms (20b).

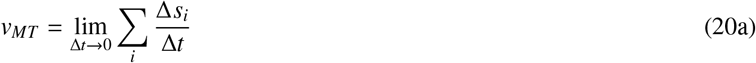

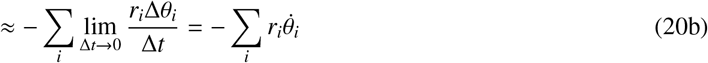

However, moment arms are not constant, and the equations for *l_MT_* and *v_MT_* must account for changes in moment arms with respect to joint angles. As a first approximation, previous work evaluated equation (20b) with posture dependent moment arms (*r_i_*(*θ_i_*) equation (21)) [18].

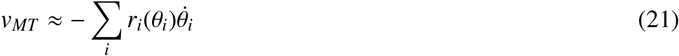

Integrating equation (21) with respect to time reveals that this approximation equates *si* to the integral of the moment arm function (*r_i_*(*θ_i_*)) across some joint rotation from neutral (*θ_i_* − *θ_i,o_*, equation (22)).

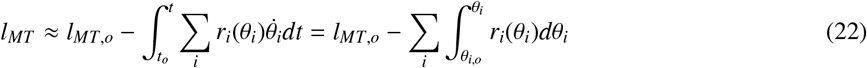

While equations (22) & (21) better approximate *l_MT_* and *v_MT_*, respectively, they fail to capture how moment arm values change *with respect to joint angles*. To capture this, we propose a new equation for *l_MT_* that relies on the definition of *arc length in polar coordinates* (23).

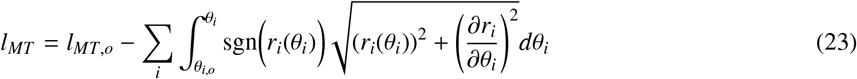

Note that when moment arms are constant, equation (19) is recovered. The sgn(*r_ij_*(*θ_j_*)) function returns +1 if *r_ij_*(*θ_j_*) > 0 and −1 if *r_ij_*(*θ_j_*) < 0 to recover the original relationship between joint rotations and excursion changes (i.e., positive joint rotation would shorten a MT with a positive moment arm). From equation (23) we derive the new equation for *v_MT_* (24b).

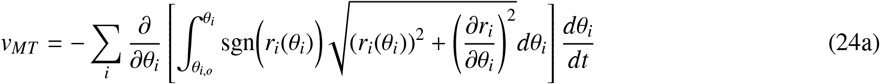

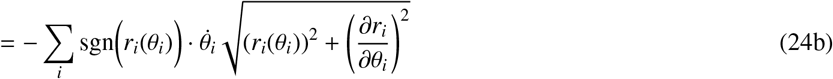

The relationship between excursions used in equations (19), (22), & (23) can be explained graphically in Figure 7 (*purple*, *orange*, and *green*, respectively).

**Figure 7:**
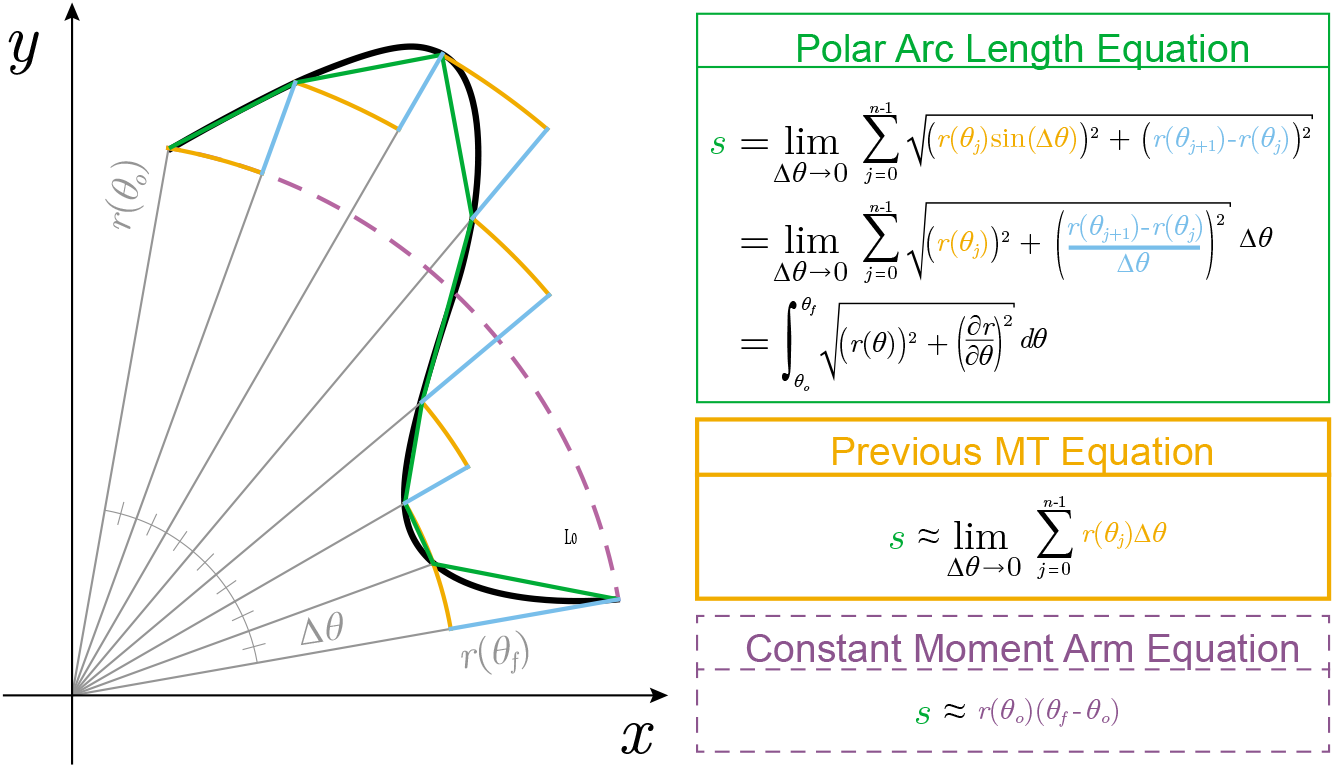
Evolution of MT excursion equations (*s*) and their differences. The constant moment arm equation [5–7] is the simplest approximation but clearly the arc length (*dashed purple*) does not accurately convey the true MT excursion (*black*). This was extended in [18] where the true arc length was approximated by integrating the posture-specific moment arm function (orange). Even for sufficiently small Δθ, this approach does not completely capture the true MT excursion as it ignores the *change in moment arm with respect to the joint angle*. Correcting for this, we find the true MT excursion from the equation for arc length in polar coordinates (*green*). As the true MT excursion relies on the Euclidean of the moment arm and its partial derivative, the error between the approximation proposed in [18] and the true equation derived here can be bounded by the triangle inequality (see equation (25)).

By exploiting the limit definition of the integral terms for MT excursion, it is easy to show that the magnitude of the error between equations (23) & (22), *ε_MT_*, for each joint will be bounded by the triangle inequality (25).

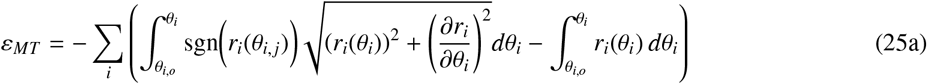

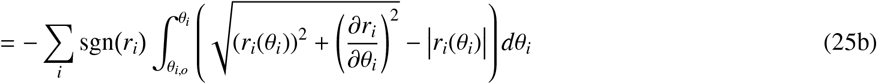

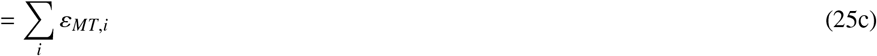

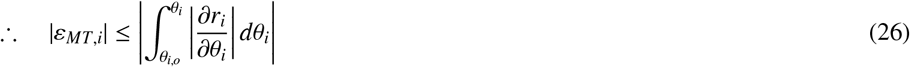

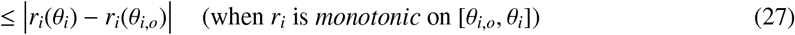

Note that the sign of the error will depend on the sign of the moment arm as well as the direction of the joint rotation such that sign of the error will be consistent with the sign of the MT behavior (i.e., *positive* for *lengthening* and *negative* for *shortening*). Therefore, errors of this type are always *underestimates* bounded between zero and equation (26) (the order of which depends on whether the joint rotation induced MT lengthening or shortening, table 1). Lastly, when *r_i_* is either increasing or decreasing *only* during the joint rotation then this error is simply bounded between zero and the difference between the moment arm function evaluated at the initial and final posture (27).

**Table 1:**
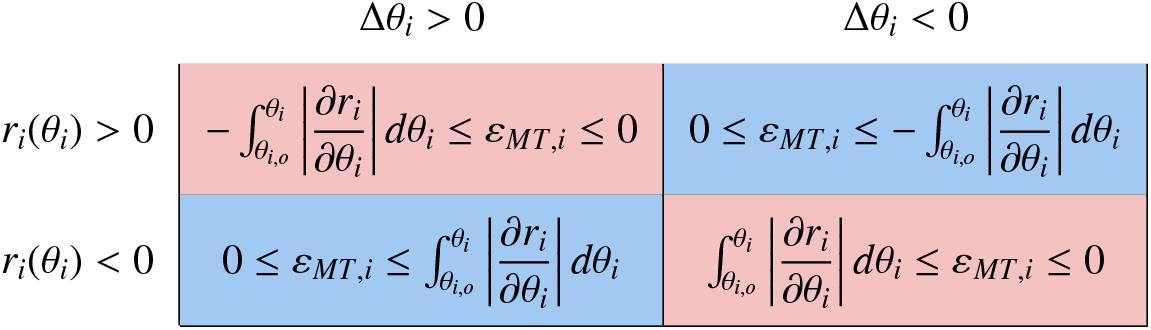
When calculating the MT excursionexcursion induced by the rotation of a joint, ignoring how much the moment arm (*r_i_*) changes with respect to the joint angle (*θ_i_*) will result in an *underestimate* of how much the MT has either *shortened* (red) or *lengthened* (blue). Whether or not a given joint rotation will caused shortening or lengthening depends on the sign of moment arm (*r_i_*(*θ_i_*)) as well as the sign of the change in joint angle (Δ*θ_i_*) [5]. Therefore, the associated error in the MT excursion (*ε_MT,i_*) will have the same sign as the MT excursion and will be bounded by the intersection of this condition and equation (26) – which both change with the signs of *r_i_*(*θ_i_*) and Δ*θ_i_*.

## A.2. Defining *c^T^* and *k^T^*

The tendon force-length relationship (given by [31], equation (7)) has fitting constants *c^T^*, *k^T^*, and 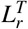. As described in [31], these parameters affect the asymptotic slope of the linear region, the curvature of the plot, and the lateral-shift of the relationship, respectively. Additionally, by restricting *c^T^* and *k^T^* to *c^T^k^T^* < 0.20, 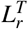 can be approximated as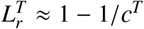 with ≈ 0.04% error in 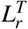, allowing for the often-unknown fitting parameter space to be constrained to 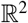. It can be shown from the limit of the slope of the *normalized* tendon force-length curve (equation (7)) as 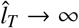 that *c^T^* is proportional to the asymptotic slope (equation (28)).

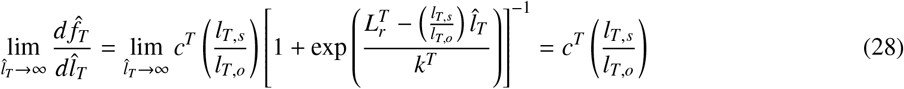

Another generalization of the tendon force-length relationship is the *stress-strain curve* where stress is given by the force normalized by the tendon’s physiological cross-sectional area (*σ* = *f_T_/CSA_T_*) and strain is given by the deformation of the tendon, normalized by the tendon’s slack length (*ε* = (*l_T_* − *l_T,s_*)/*l_T,s_*) [3, 29, 51, 56]. This representation is useful for calculating the elastic modulus of tendon (*E_T_*, Young’s modulus for tendinous tissue) as the slope of the linear region.

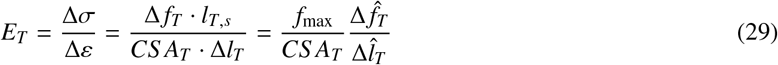

As the slope of the linear region of (7) is given by *c^T^*(*l_T,s_/l_T,o_*), we can rewrite *c^T^* as (30). This equation expands upon the definition of *c^T^* as the “linear stiffness” parameter and provides physical intuition about how changes in physiological parameters like tendon cross sectional area and muscle force producing capabilities will affect it.

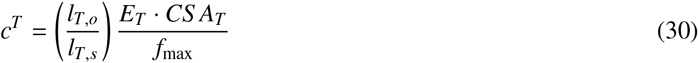

As the result of this relationship, tendon length changes at high forces can be approximated as

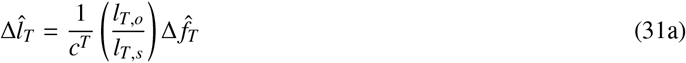

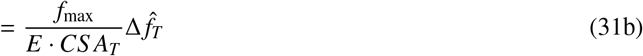

How the fitting parameter *k^T^* relates to the curvature of “toe” region of the normalized tendon force-length relationship is not explicitly clear. To explore the effect that changes in these parameters have on curvature, we define curvature as (32). This definition considers the amount of change that the tangent vector along a curve has in the direction of the normal vector. In the case where the curve is a circle, the curvature is defined as the inverse of the radius. Thus, as the radius *decreases*, the curvature *increases* and the “sharpness” of the curve increases. In the case where the radius becomes very large, the curvatures goes to zero as the circle locally approaches a straight line. Therefore, large curvature values (*κ*) are associated with sharp changes along the curve. In mechanical systems, this is often approximated as the second derivative of the curve.

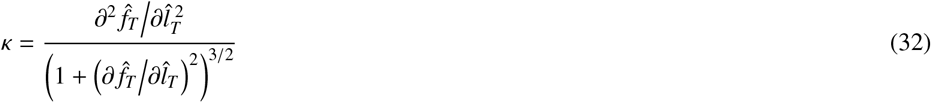

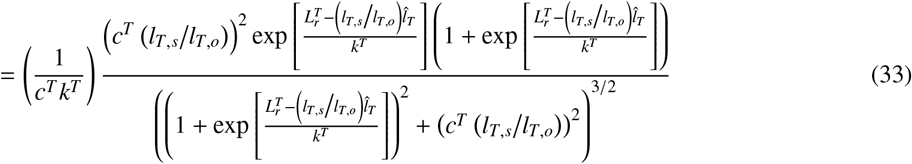

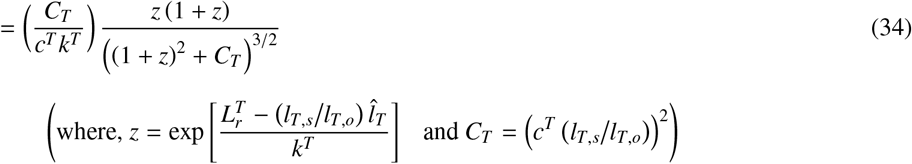

As *k^T^* is defined as the variable that affects curvature (which varies along the continuous curve), we derive the maximum curvature of the tendon force-length curve in order to see the influence that *k^T^* has on it. To do so, we take the derivative of (32) with respect to 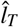 and find its zeros.

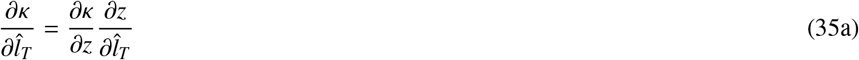

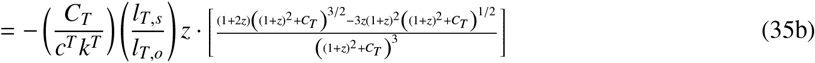

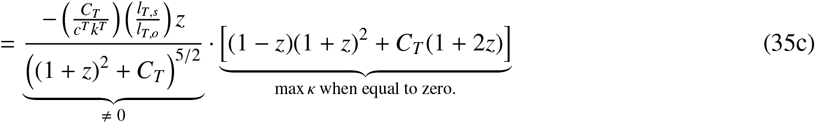

Therefore the curvature is at a maximum when

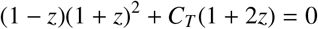

which will occur when,

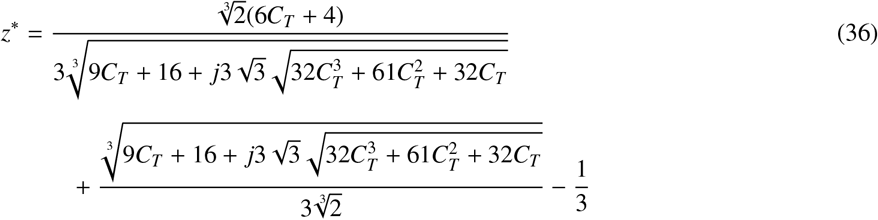

Representing (36) in terms of magnitude and phase allows us to remove the imaginary component.

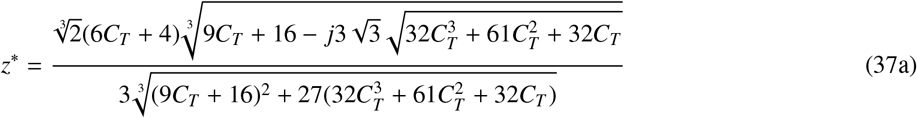

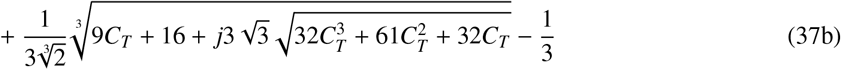

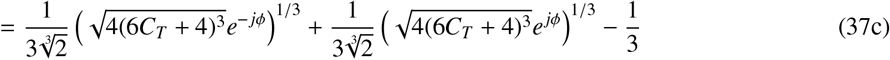

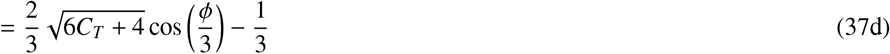

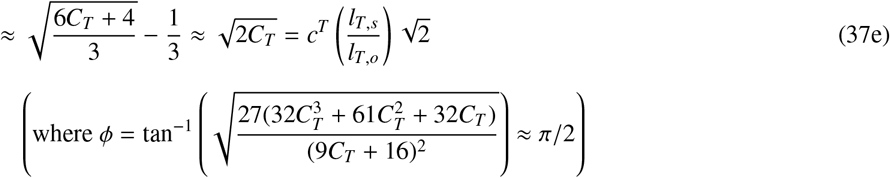

Therefore the maximum curvature is given by plugging the (37c) into (34).

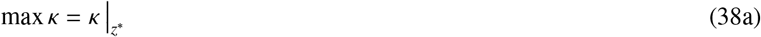

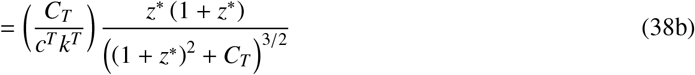

The approximate maximum curvature is found by instead evaluating the curvature at the approximate value for *z*\* (equation (37e)). If we make the reasonable assumption that *c^T^* >> 10 (see *Appendix A.3*), we can manipulate the equation for maximal *κ* to find,

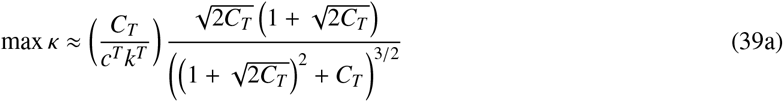

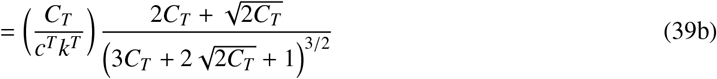

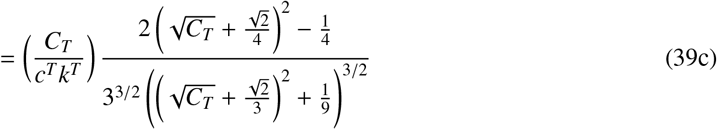

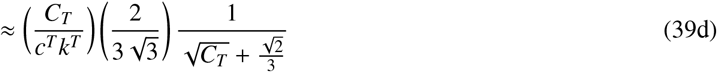

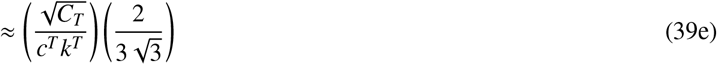

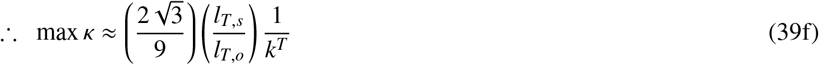

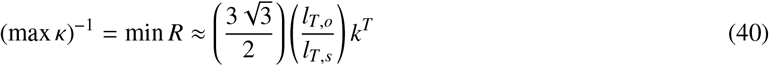

Therefore, as equation (39f) suggests, the value of *k^T^* is *inversely* proportional to the curvature and therefore proportional to the *radius* of curvature of the “toe” region (R), where small values of *k^T^* correspond to high curvature or low radius of curvature (i.e., would exhibit *sharp transitions* from the “toe” region to the “linear” region). Additionally, equation (39f) and its reciprocal, equation (40), could be used to help find the hard-to-measure *k^T^* constant from an experimental, normalized tendon force-length curve by either calculating the curvature and finding its maximum or by measuring the smallest radius of curvature, respectively.

## A.3. Defining Physiological Ranges for *c^T^* & *k^T^*

The parameters that are used to characterize the *normalized* tendon force-length curve—asymptotic stiffness, the radius of curvature constant, and lateral shift (*c^T^, k^T^*, and 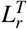, respectively)—greatly influence the behavior of the tendon and vary across muscles, subjects, and sometimes tension rates [57, 58]. In order to determine the effects that changing these fitting parameters have on overall MT behavior, we define the physiological range for *c^T^* and *k^T^* from (i) the condition that *c^T^k^T^* < 0.20 [31], and (ii) the condition that, by definition, the tendon must be at its slack length 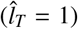 when normalized tendon force 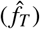 is near zero [3, 31, 32, 51]. The first constraint on *c^T^* and *k^T^* comes from [31], who stated that restricting values to *c^T^k^T^* < 0.20 allows 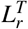 in equation (7) to be approximated as 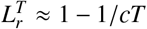 while incurring only a 0.04% error in 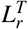 (*red* region excluded in figure 8). The second constraint produces a range of acceptable values of *c^T^* and *k^T^* when satisfying equation 8 for 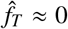 and 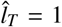 and for *l_T,s_/l_T,o_* ∈ [1.03,1.0.7] (*blue* regions excluded in figure 8) [51, 52].

**Figure 8:**
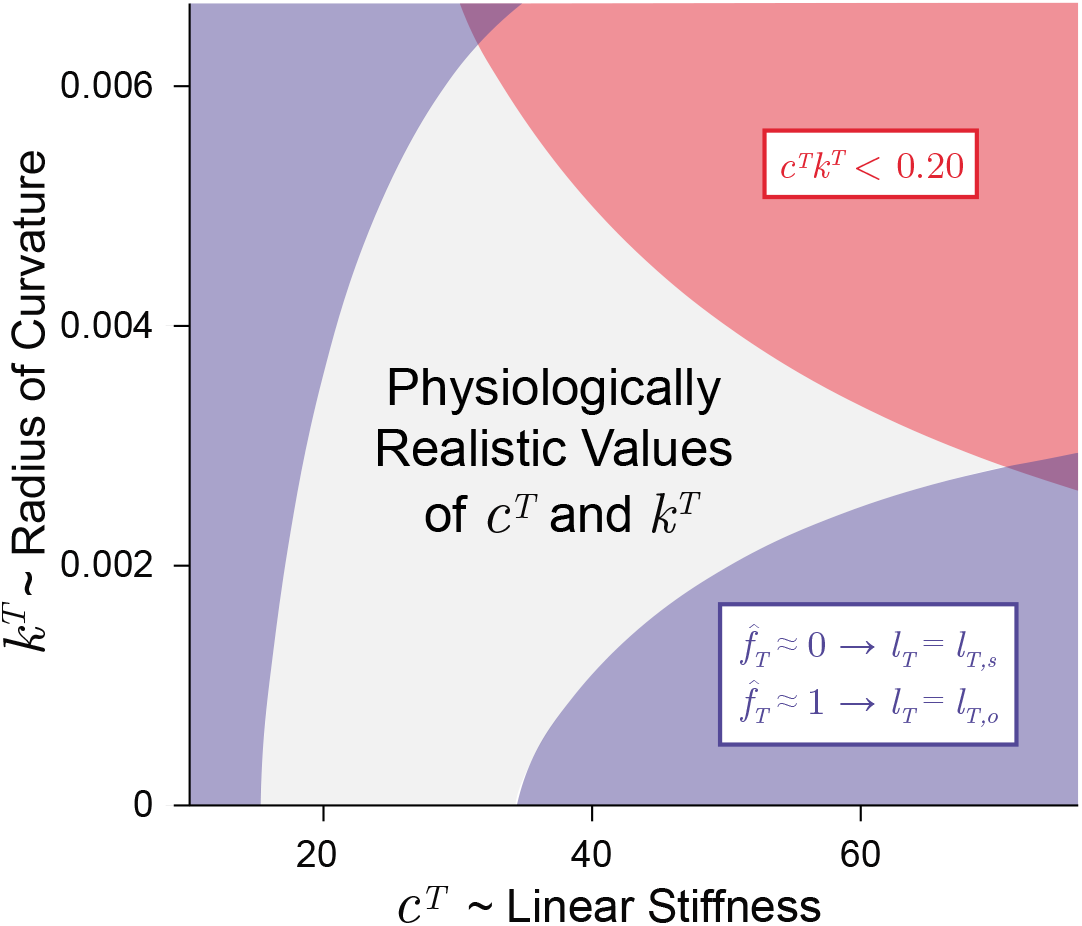
Physiologically realistic ranges for *c^T^* and *k^T^* under the assumptions that (i) *c^T^k^T^* < 0.20 (*red*) [31], and (ii), by definition, when force is negligible, tendon length equals its slack length (i.e., 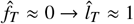, *blue*) [3, 31, 32, 51].

To validate the ranges produced by these constraints, we explore the reported ranges for *c^T^* as (i) we can calculated *c^T^* from Young’s modulus (E), the tendon’s physiological cross-sectional area (*CSA_T_*), and the maximum isometric force of the muscle (*f_max_*) from equation 30 and (ii) the values of *k^T^* are less often reported. Young’s modulus has been reported to be conserved across muscles with the average *E* reported to be around 1.2 GPa and, therefore, changes to *c^T^* can be attributed to changes in *CSA_T_* and *f_max_* [3, 51, 59, 60]. These two parameters change across MTs as well as with training, injury, or pathology and help to explain the large variability in tendon stiffness seen across muscle and subjects [52, 61, 62]. As an example, Magnusson *et al*. calculated E, *CSA_T_*, and *f_max_* for the *medial gastrocnemius* of 5 individuals during isometric contraction tasks [51] and the *c^T^* values calculated from equation (30) ranged from 23.23 to 65.70 (37.47 ± 14.88). Therefore, the range of *c^T^* values produced by the two constraints described above are consistent with values reported in the literature and are good first approximations of the range of physiological values when exploring the affect they have on tension-specific tendon deformation.

## Footnotes

1 I.e., by tracing the MT routing from its origin to insertion across postures (See [4]), or by calculating the integral of the moment arm functions for all joints crossed (See [5–7]). The latter approach does not consider how moment arms change with *respect to joint angles*. In *Appendix A.1*, we derive more accurate equations for MT length and velocity that account for this.

2 While these two assumptions are typically made together, they have been separated here to demonstrate the relative error contribution for which each assumption accounts.

3 See *Appendix A.2 & A.3* for more in depth description of these parameters and their physiological ranges.

4 Assuming that *rate*-specific phenomenon like *hysteresis, creep, force-relaxation* and *short-range stiffness* are negligible.

5 *C*_1_ sensitivity can also be defined as the partial derivative of *C*_1_ evaluated at *p_c_*. 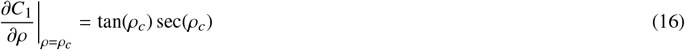 Note that as *ρ_c_* increases, so does the slope of *C*_1_ evaluated at *ρ_c_*, indicating that for the same small deviation from p_c_, the proportion of Δ*l_MT_* that was incorrectly projected back onto the line of action of the fibers will increase as *ρ_c_* increases.

6 *C*_2_ sensitivity can also be defined as the partial derivative of *C*_2_ evaluated at *ρ*(*t_o_*). 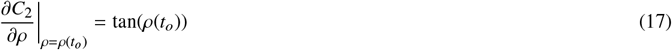 Similar to *C*_1_, as *ρ*(*t_o_*) increases, so to does the slope of *C*_2_ evaluated at *ρ*(*t_o_*), indicating that for the same small deviation from*ρ*(*t_o_*), the proportion of *l_m_*(*t_o_*) that was incorrectly projected back onto the muscle will increase as *ρ*(*t_o_*) increases.

7 Given that *l_m_*(*t_o_*) = 5.6 cm and *l_m,o_* = 4.8cm [44]

8 The range for *ρ* was chosen to be < 40° (See *Section 3.1*), the range for *l_T,o_/l_T,s_* was chosen to be 1.03-1.07 [51, 52, 54], and the range for *l_T,s_/l_m,o_* has been reported to be ≤ 11.25 [3, 55]. For an explanation of the physiological levels of *c^T^* and *k^T^* see *Appendix A.3*.

## Notes

https://daniel8hagen.com/images/tendon_length_change_parallel_coords

### Competing Interest Statement

The authors have declared no competing interest.

